# ADAT2/3-mediated tRNA editing promotes cancer cell growth and tumorigenicity

**DOI:** 10.1101/2024.10.31.621298

**Authors:** Julia Ramirez-Moya, Titi Rindi Antika, Qi Liu, Xushen Xiong, Raja Ali, Alejandro Gutierrez, Richard I. Gregory

**Affiliations:** Stem Cell Program, Boston Children’s Hospital, Boston, MA 02115, USA; Division of Hematology/Oncology, Boston Children’s Hospital, Boston, MA 02115, USA; Department of Biological Chemistry and Molecular Pharmacology, Harvard Medical School, Boston, MA 02115, USA; Rice Research Institute, Guangdong Academy of Agricultural Sciences, Guangzhou 510640, Guangdong Province, China; Guangdong Key Laboratory of New Technology in Rice Breeding, Guangzhou 510640, Guangdong Province, China; Zhejiang University Medical Center, Hangzhou, Zhejiang Province, China; Department of Pediatrics, Harvard Medical School, Boston, MA 02115, USA; Harvard Initiative for RNA Medicine, Boston, MA 02115, USA

**Author notes:** Corresponding author: Richard I. Gregory.

**Keywords:** Editing, Inosine, ADAT2, ADAT3, tRNA, epitranscriptome, mRNA translation, cancer

## Abstract

Transfer RNAs (tRNAs) are subject to various chemical modifications that influence their stability or function. Adenosine to Inosine (A-to-I) editing in the tRNA anticodon at position A34 is an important modification that expands anticodon-codon recognition at the wobble position and is required for normal mRNA translation. The relevance of tRNA editing in cancer remains unexplored. Here we show that the genes encoding the ADAT2/3 deaminase complex, responsible for A-to-I tRNA editing in humans, are commonly amplified and/or overexpressed in several tumor types including liposarcoma (LPS). We find that knockdown of the ADAT complex suppresses LPS cell growth and tumorigenicity. Mechanistically, we find that decreased tRNA editing upon ADAT2 depletion leads to defective translation of a subset of mRNAs. Thus, ADAT-mediated tRNA modification promotes oncogenesis by enhancing the translation of growth promoting mRNAs that are enriched in NNC codons that lack cognate tRNAs and therefore depend on A-I tRNA editing for decoding and mRNA translation. Our results uncover an oncogenic role of tRNA editing and identify ADAT2/3 as a potential new cancer therapeutic target.

---

Transfer RNAs (tRNAs) are key adaptor molecules essential for messenger RNA (mRNA) translation and are subject of numerous posttranscriptional modifications, including Adenosine-to-Inosine (A-to-I) editing, the major RNA editing type that occurs in humans[1]. The enzyme responsible for this modification is the Adenosine Deaminase Acting on tRNA (ADAT) complex comprising the catalytic ADAT2 component and ADAT3 co-factor that is essential for recognition and presentation of the tRNA to the catalytic pocket for editing the subset of human tRNAs with adenosine at position 34 (A34)[2], [3], [4], [5], [6]. tRNA nucleotide 34 is located within the anticodon, which faces the third base of mRNA codons during translation. Modification of A-to-I at this wobble position has an obvious implication for anticodon:codon recognition because, while A34 can base-pair with U, I34 is capable of pairing with A, U, and C (**Figure 1A**). This expands the repertoire of triplets that the modified tRNA can recognize and, in doing so, profoundly modifies the balance between codon usage and tRNA abundance. Although only tRNA-Arg-ACG is modified in most prokaryotes, up to eight tRNAs are inosine-modified in eukaryotes including tRNA-Ala-AGC, tRNA-Arg-ACG, tRNA-Ile-AAT, tRNA-Leu-AAG, tRNA-Pro-AGG, tRNA-Ser-AGA, tRNA-Thr-AGT and tRNA-Val-AAC[2]. In fact, eukaryotic ADAT emergence was accompanied by a dramatic genomic expansion of A34-tRNA genes, and most eukaryotic genomes lack tRNA genes that produce G34-tRNAs[7], [8]. Therefore, A-to-I editing is required to compensate for the lack of G34-tRNAs otherwise needed to decode C-ending (NNC) codons[9]. Mutation in the *Schizosaccharomyces pombe tad3* gene (ortholog of the human *ADAT3* gene) causes diminished deaminase activity of the complex and results in defective cell cycle progression[10]. ADAT2 or ADAT3 knockdown in human HEK293T cells similarly leads to defects in cell cycle progression as well as cell adhesion phenotypes[11]. Germline mutations in *ADAT3* resulting in compromised A-I tRNA editing are associated with intellectual disability in human patients[12], [13], [14], and conditional *Adat2* deletion in the B cell lineage causes a near complete loss of peripheral blood B cells in mice[15].

**Figure 1.**
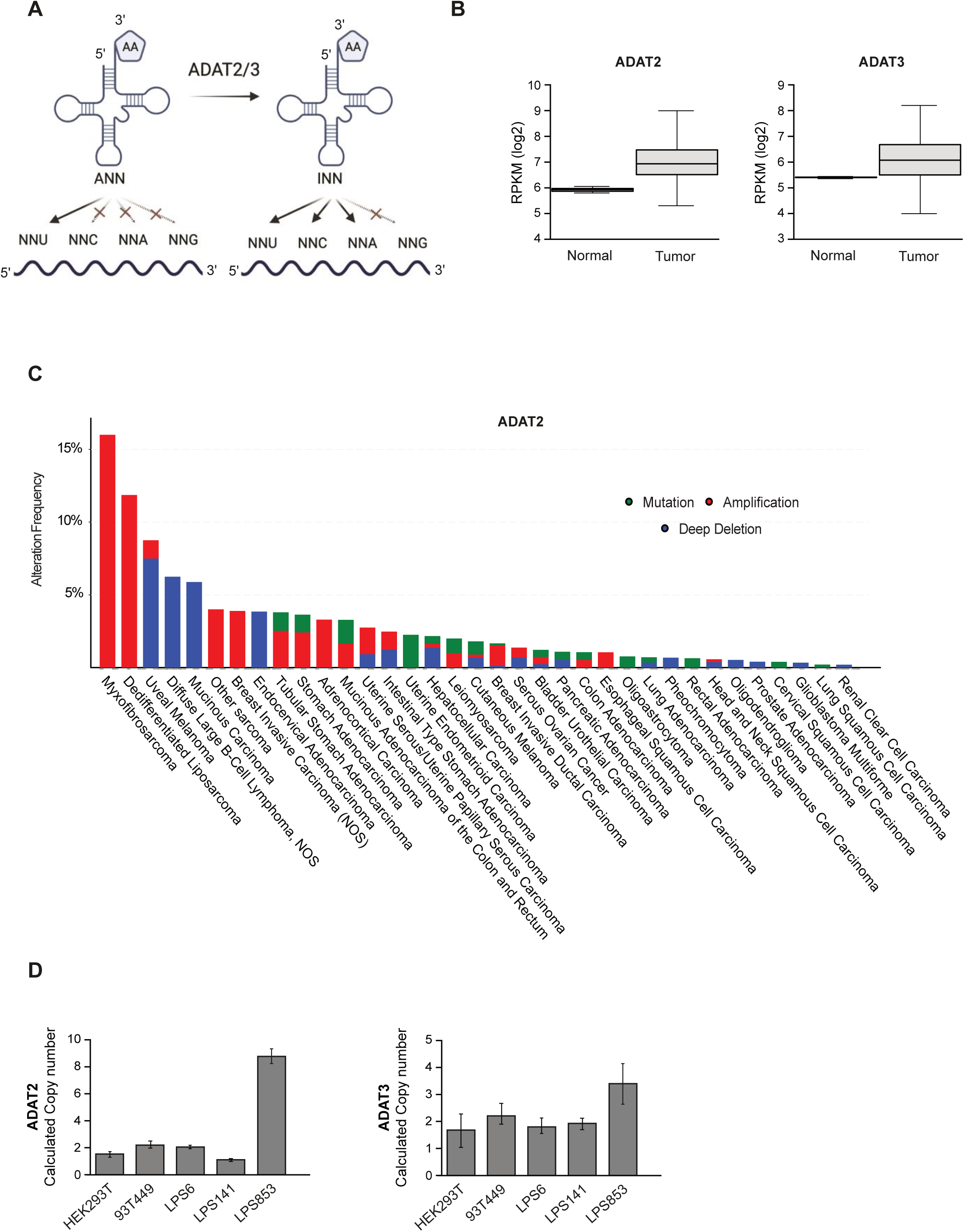
*ADAT2/3* are amplified and overexpressed in liposarcoma tumors. (**A**) Schematic representation of A-to-I tRNA editing and its role in codon recognition. (**B**) ADAT2 (left) and ADAT3 (right) mRNA levels in sarcoma tumors and corresponding normal samples from the TCGA database. (**C**) *ADAT2* genetic alterations (mutations, amplifications, and deletions) in TCGA tumors. ‘Other sarcomas’ refers to Undifferentiated Pleomorphic Sarcoma/Malignant Fibrous Histiocytoma/High-Grade Spindle Cell Sarcoma. (**D**) *ADAT2* and *ADAT3* copy number in the indicated liposarcoma cell lines.

Codon composition is an important factor that influences translation rates, and the clustering of codons in mRNAs that have a low level of cognate tRNAs limits the rate of translation[16]. Thus, the translation of genes rich in ADAT-sensitive codons (codons translated by I34-tRNAs) might benefit from the increased decoding capacity of inosine-modified tRNAs. In agreement with this prediction, self-renewing mouse and human embryonic stem cells (ESCs) that express many genes enriched in ADAT-sensitive codons display elevated I34 levels[17].

Here we explore for the first time the relevance of A-to-I editing of tRNAs in cancer. Analysis of The Cancer Genome Atlas (TCGA) (https://www.cancer.gov/tcga), identified that the components of the ADAT complex are dysregulated in cancer. We observed that sarcoma tumors, and specifically liposarcomas, are the cancer types that present higher genetic alterations in *ADAT2* and *ADAT3*. Liposarcoma (LPS) is a rare cancer that arises in fat cells, typically in the soft tissue of the body. There are several subtypes of liposarcoma, each with different clinical and pathological features. The most common subtype is well-differentiated liposarcoma, which is typically slow-growing and has a low propensity for metastasis. Other subtypes include myxoid liposarcoma, pleomorphic liposarcoma, and dedifferentiated liposarcoma, which are generally more aggressive and have a higher risk of spreading to other parts of the body[18], [19]. The primary treatment for liposarcoma is surgical removal of the tumor, with radiation therapy or chemotherapy used in some cases[20], [21]. However, some tumors present a high risk of metastasis and a poor prognosis, thereby highlighting the importance of identifying new molecular markers for better diagnosis and possible future therapeutic targets.

Here we demonstrate that the ADAT complex has an oncogenic role in LPS. We show that increased tRNA editing in cancer cells promotes the translation of oncogenic mRNAs that are enriched in ADAT-dependent codons, providing a selective advantage to cancer cells. Our data provide new insights into the role of A-to-I tRNA editing in gene expression and cancer and we propose that the ADAT2/3 deaminase could represent a new therapeutic target for cancer treatment.

## Results

### *ADAT2* and *ADAT3* are amplified and overexpressed in human tumors

To explore the possible altered mRNA expression levels of ADAT2 and ADAT3 in cancer we analyzed available RNA-Seq data from The Cancer Genome Atlas (TCGA) (https://www.cancer.gov/tcga). This revealed elevated ADAT2 and ADAT3 mRNA expression levels in most types of human tumors compared to corresponding normal tissues (**Figure S1A-B**). Moreover, gene copy number analysis showed that *ADAT2* and *ADAT3* loci are frequently amplified in several tumor types **(Figure 1C**, **Figure S2**). Taken together, the cancer genetics and mRNA expression data implicate ADAT2 and ADAT3 in various cancers. More detailed examination of TCGA datasets revealed that ADAT2 and ADAT3 mRNA expression is elevated in sarcomas compared with normal tissue samples (**Figure 1B**). Among the sarcoma group, frequent amplification of *ADAT2* and *ADAT3* genes is observed in the dedifferentiated liposarcoma (LPS) subset (**Figure 1C, Figure S2**). Therefore, we considered LPS a relevant model in which to study the role of ADAT proteins in cancer and to explore the underlying molecular mechanisms.

### ADAT2 and ADAT3 are required for cancer cell growth and tumorigenicity

To explore the requirement for ADAT2/3 in cancer cells, we used short harpin RNA (shRNA) constructs to specifically knock down ADAT2 and ADAT3 in human LPS cell lines. We used two different shRNAs in two LPS cell lines with different amplification levels of ADAT2 and ADAT3. According to our results, the LPS853 cell line harbors nine copies of *ADAT2* and four copies of *ADAT3* (**Figure 1D**). Considering this, we used LPS853 as a model LPS cell line with *ADAT2/3* gene amplification and the LPS6 cell line as a non-amplified LPS cell line (**Figure 1D**). These two cellular models allowed us to examine the requirement of ADAT2/3 in human LPS cell lines and to furthermore determine if the cells with a higher amplification and/or expression of ADAT2/3 are more sensitive to its silencing.

We evaluated the effects of ADAT2 depletion on cancer cell growth and viability using a panel of cellular assays to measure several hallmarks of cancer and comparing our knockdown LPS cells to the parental cell line or cells transduced with a non-targeting shRNA (shGFP) as controls (**Figure S3**). We observed that depletion of ADAT2 caused decreased cell proliferation (**Figure 2A**), colony formation (**Figure 2B**), invasion (**Figure 2C**), and anchorage-independent cell growth (**Figure 2D**). Interestingly, we found that ADAT2-amplified LPS853 cells are more sensitive to ADAT-depletion than non-amplified LPS6 cells. Moreover, in contrast to the LPS cancer cell lines, knockdown of ADAT2 in the non-cancerous human BJ fibroblasts did not cause any strong growth defects (**Figure S4A-B**). These results suggest that ADAT2 promotes cancer cell growth and is essential for maintaining a malignant phenotype in LPS cell lines. To further explore this hypothesis, we assessed the ability of the stable ADAT2 KD cell lines to form tumors *in vivo* in immunocompromised mice. Strikingly, we observed that silencing of this enzyme strongly suppressed the ability of these cells to form tumors in these xenograft studies (**Figure 2E**). These results highlight ADAT2 as a possible new therapeutic target in cancer. Very similar results were observed upon ADAT3 silencing (**Figure S5A, B**), with ADAT3-depleted LPS cells showing decreased proliferation (**Figure S5C**), colony formation (**Figure S5D**), invasion (**Figure S5E**), and anchorage-independent cell growth (**Figure S5F**), which is consistent with the ADAT3 subunit being required for tRNA editing activity of the ADAT2/3 complex. As such, ADAT3 might also be also considered as a potential anti-cancer therapeutic target.

**Figure 2.**
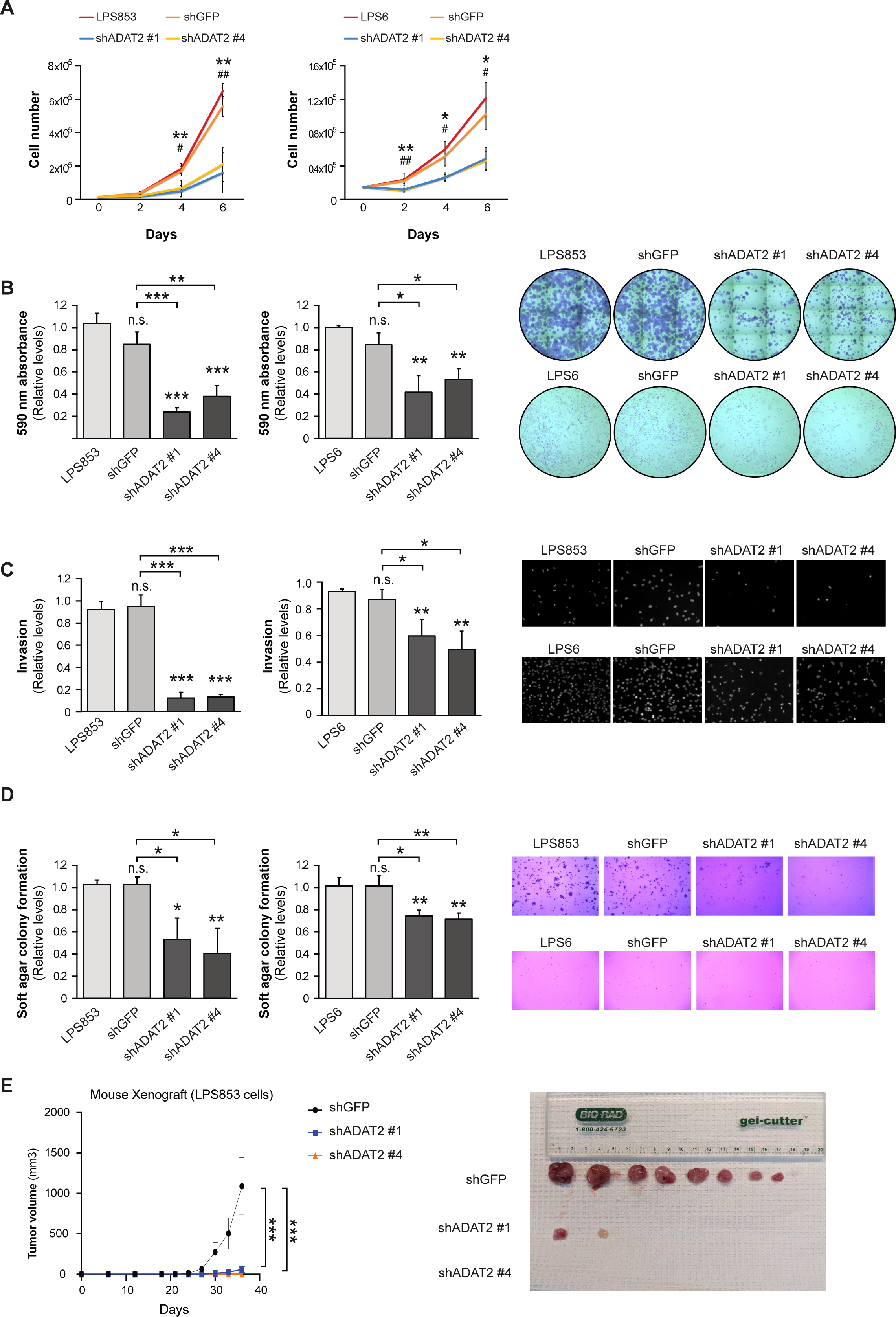
ADAT2 is required for liposarcoma cell growth and tumorigenicity. ADAT2-knockdown in human liposarcoma (LPS) cell lines, LPS853 and LPS6, cells using two different shRNAs (shADAT2 #1 and shADAT2 #4). shGFP and wild-type cells were used as negative controls. (**A**) Proliferation assay (n=3). (**B**) 2D colony formation assay. Quantification (left) and representative images (right) (n=3). (**C**) Invasion assay. Quantification (left) and representative images (right) (n=3). (**D**) 3D soft agar colony formation assay. Quantification (left) and representative images (right) (n=3). (**E**) Tumor growth in mouse xenografts using LPS853 cells. Quantification (left) and tumor images at endpoint (right). Error bars indicate standard deviations. Asterisks denote statistical significance assessed with Student’s t-test (two-tailed). * p < 0.05, ** p < 0.01, *** p < 0.001.

### ADAT2 catalytic activity promotes growth of liposarcoma cells

To ensure that the growth phenotypes observed in LPS cells are due specifically to ADAT2 deficiency and to rule out possible off-target effects of the shRNA, we performed rescue assays in which an ADAT2 overexpression vector was introduced into the ADAT2 KD cells (**Figure 3A-B**) For this purpose, we generated an overexpression vector containing the coding sequence of ADAT2 and we performed directed mutagenesis to introduce silent mutations within the open reading frame (ORF) to avoid the specific silencing by the ADAT2 shRNA previously employed. We repeated the above-mentioned functional assays to compare the KD cells to the rescued ADAT2 in the shADAT2 condition and the control shGFP cells. Our results showed that the observed phenotypes are specific for the ADAT2 KD, as the recovery of ADAT2 expression rescues the proliferation, invasion, and soft agar colony formation capability in both LPS853 and LPS6 cell lines (**Figure 3C-E**). In addition, we also studied the importance of ADAT2 catalytic activity by including a catalytic mutant version of ADAT2 (ADAT2 E73A)[2] in our rescue experiments. Our results showed that only the wild-type ADAT2 and not the catalytically inactive mutant form is able to rescue the malignant phenotype (**Figure 3C-E**), therefore strongly supporting that the tRNA-editing activity of ADAT2 is responsible for the observed growth effects.

**Figure 3.**
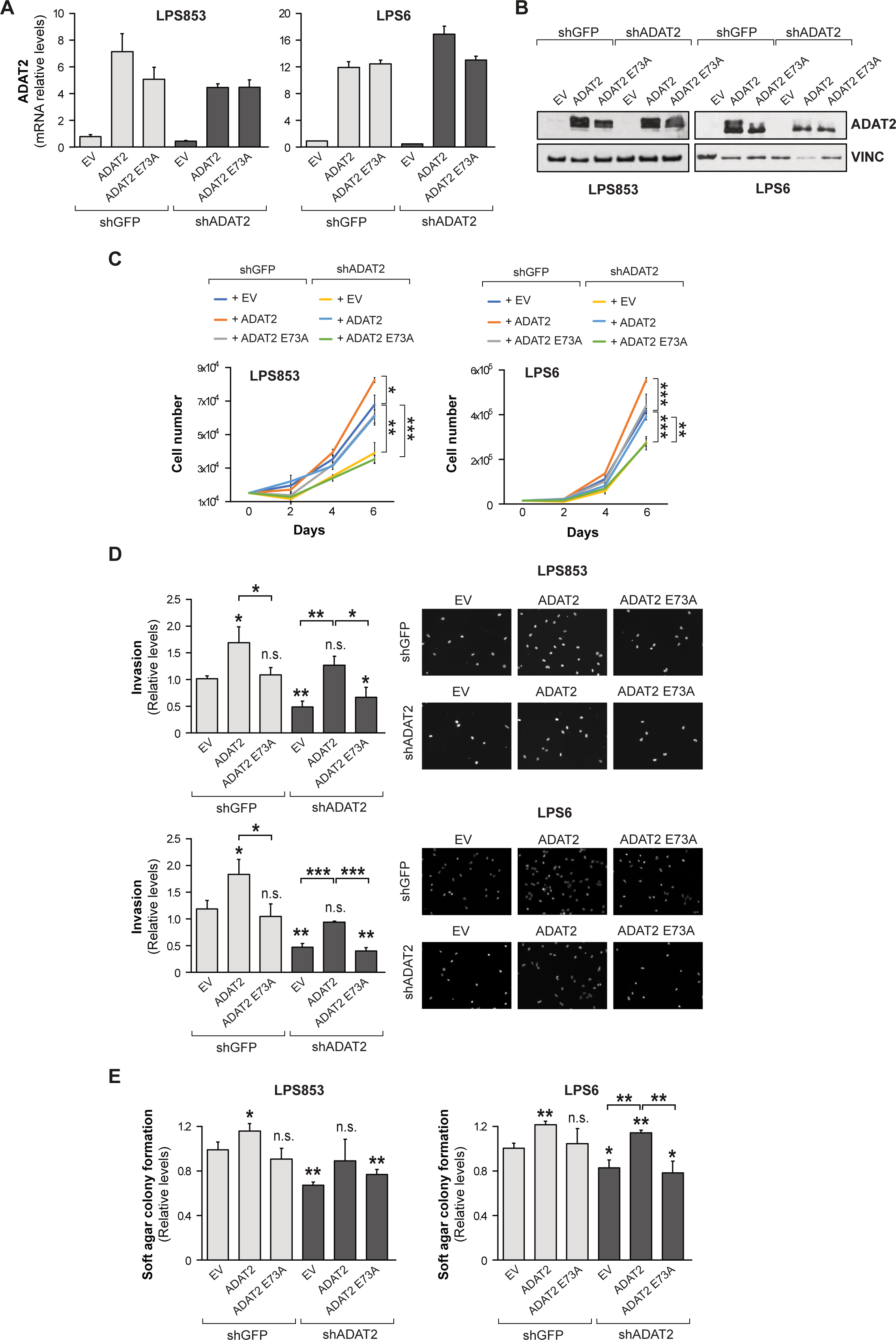
ADAT2 catalytic activity promotes liposarcoma cell growth. LPS853 and LPS6 cells were transduced with an ADAT2 shRNA (shADAT2) or with shGFP as a control. Then, ADAT2 wild-type or the catalytic mutant ADAT2 E73A were overexpressed to perform rescue assays. (**A**) Relative ADAT2 mRNA levels assayed by qPCR (n=3). (**B**) Representative Western Blotting for ADAT2 protein. Vinculin (VINC) was used as loading control. (**C**) Proliferation assay (n=3). (**D**) Invasion assay. Quantification (left) and representative images (right) (n=3). (**E**) 3D soft agar colony formation assay (n=3). Error bars indicate standard deviations. Asterisks denote statistical significance assessed with Student’s t-test (two-tailed). * p < 0.05, ** p < 0.01, *** p < 0.001.

### Decreased A-to-I tRNA editing in ADAT2-depleted LPS cells

To investigate the molecular mechanism involved in the oncogenic role of the ADAT2/3 complex, we first performed small RNA sequencing in the LPS853 control and ADAT2-knockdown cells and measured A-to-I tRNA editing levels. Inosine is resolved as guanosine upon cDNA sequencing. Consistent with the previous report on HEK239T cells[22], the proportion of inosine at position 34 is reduced by up to ∼4-fold in several tRNA isodecoder families including tRNA-Ser-AGA, tRNA-Val-AAC, tRNA-Pro-AGG, and tRNA-Thr-AGT. Other tRNAs including tRNA-Arg-ACG and tRNA-Ile-AAT, although belong to tRNA isodecoder families that are edited[23], were largely unaffected by knockdown of ADAT2 (**Figure 4**). Our results show an overall decrease in the editing of several tRNA families with some more affected than others under conditions of partial ADAT2 depletion in LPS cells.

**Figure 4.**
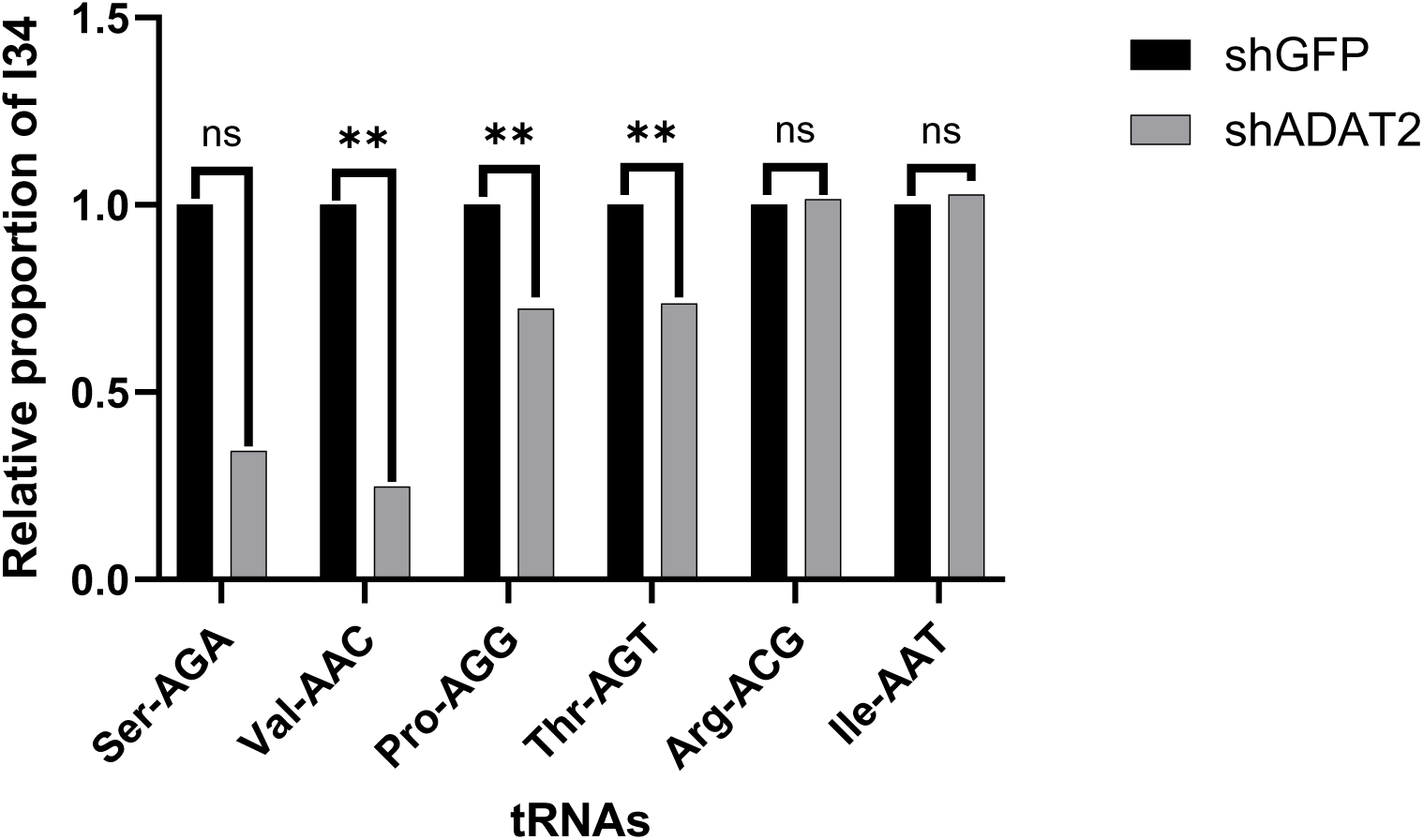
ADAT2-knockdown decreases A-to-I tRNA editing in liposarcoma cells. Relative proportion of inosine at position 34 in human ANN tRNAs observed in shGFP and shADAT2 LPS853 liposarcoma (LPS) cells (n=3), with ** p < 0.01.

### ADAT2 deficiency leads to codon-biased changes in mRNA translation

To examine the impact of ADAT2 silencing and decreased A-to-I tRNA editing on mRNA translation, we performed ribosome footprinting (Ribo-Seq) experiments in control and ADAT2-depleted LPS853 cells. Ribo-Seq revealed a remodeling of the mRNA ‘translatome’ due to ADAT2-deficiency with 183 mRNAs >1.5-fold more efficiently translated (upregulated) and 146 mRNAs less efficiently translated (downregulated) in ADAT2-knockdown compared with control cells (**Figure 5A, B, and Supplementary Dataset 1**). Interestingly, gene ontology (GO) analysis showed that the genes with decreased translation efficiency in ADAT2-depleted cells are enriched in cancer-related processes including cell migration, cell proliferation, and cell motility (**Figure 5C**). These molecular changes in the translation efficiency of growth-promoting genes likely help explain the oncogenic role of ADAT2 and the cellular growth phenotypes we observed ADAT2-depleted LPS cells (**Figures 2 and 3**). We next explored whether the mRNAs with altered translation in ADAT2-depleted cells are enriched in codons that are normally decoded by the subset of A-to-I edited tRNAs. For this purpose, we compared the codon usage between genes with increased translation efficiency (TE up) versus decreased translation efficiency (TE down) with the enrichment of codons in the TE Down subset compared with the global average codon usage across all human open reading frames (ORFs) throughout the genome. We found that several ADAT-sensitive codons are both enriched in mRNAs with decreased versus increased TE and are enriched in the TE down group compared with global average codon usage (**Figure 5D**). A significant enrichment of multiple ADAT-sensitive codons including GCC, CCC, CCA, ACC, CGC, GTC, ATC, and CGA in the mRNAs with decreased translation efficiency (TE down) upon ADAT2-knockdown was observed (**Figure 5D**). Interestingly, we found NNC codons to be the most dependent on A-to-I tRNA editing, which correlates with the general absence of G34 tRNAs in the human genome[9]. Therefore, decoding of NNC is dependent on A-I editing since these codons would otherwise lack cognate tRNAs for translation. Amongst this group are Arg-CGC, Pro-CCC, Ala-GCC, and Thr-ACC codons that are all significantly enriched in the mRNAs with decreased translation due to ADAT2 deficiency (**Figure 5E**). Conversely, NNT codons that can be decoded by both the unedited (A34) as well as by the edited tRNAs including CTT, ATT, GTT, and GCT codons have a lower dependance on tRNA editing and are thus less enriched in the mRNAs with decreased translation efficiency in the ADAT2-depleted cells (**Figure 5D**). Interestingly, the highly enriched NNC codons (GTC, CCC, and ACC) are recognized by the edited form of tRNA-Val-AAC, tRNA-Pro-AGG and tRNA-Thr-AGT, respectively, further supporting the importance of ADAT2/3-mediated editing of this subset of tRNAs for efficient translation of mRNAs containing these codons. In conclusion, ADAT2-deficiency causes selectively decreased translation of mRNAs that are enriched in NNC codons that depend on A-I tRNA editing for decoding. Thus, ADAT2-mediated tRNA editing is required for codon-biased mRNA translation of growth-promoting genes in human cancer cells.

**Figure 5.**
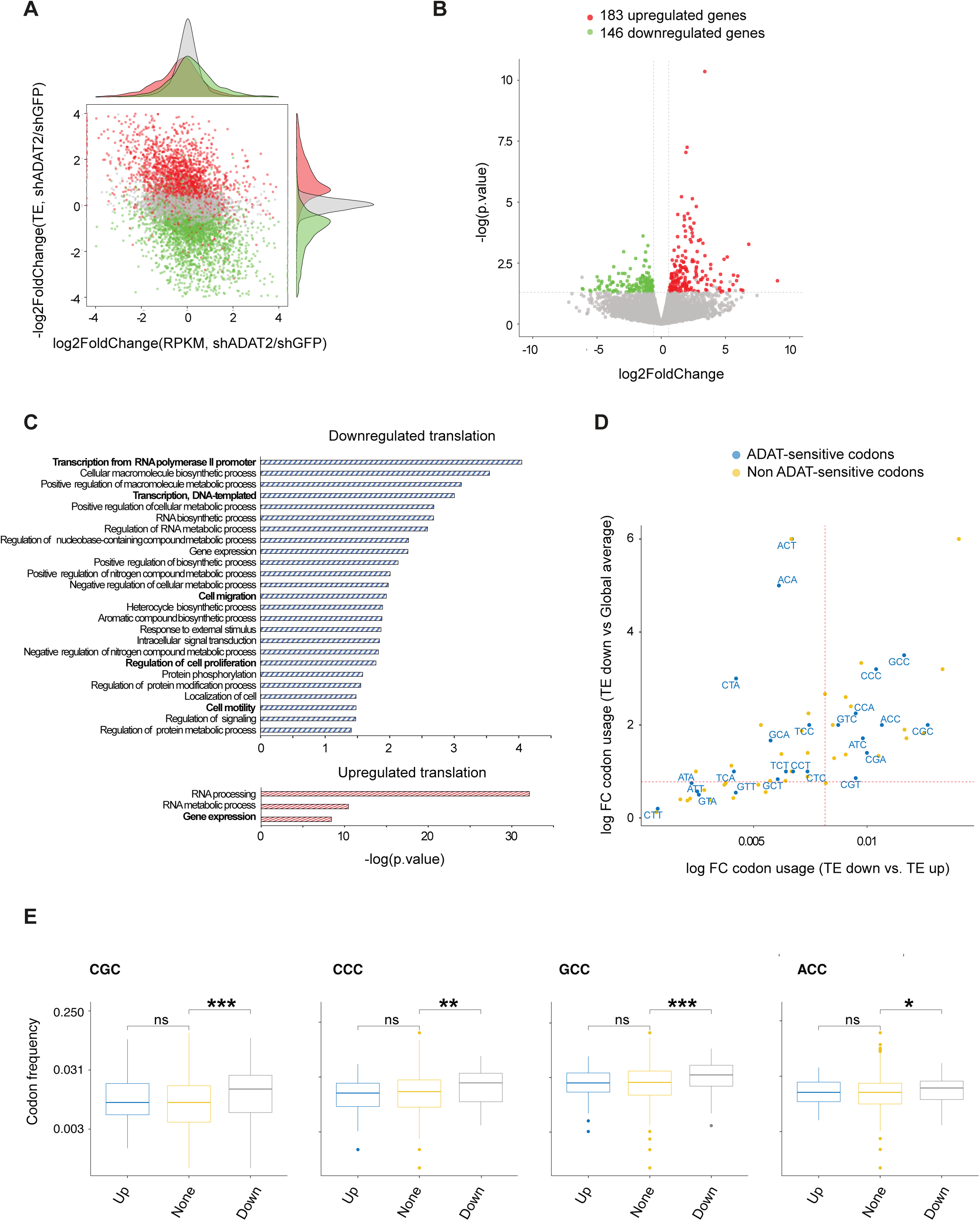
ADAT2 is required for translation of mRNAs enriched in ADAT-sensitive codons. Ribo-seq was performed in shGFP and shADAT2 LPS853 cells (n=2). (**A**) Translation efficiency (TE) change v.s. gene expression changes (red: up-translation, green: down-translation). TE was calculated by dividing the ribosome-protected fragments (RPF) signals by the input RNA-seq signals. (**B**) Translation changes observed after ADAT2 silencing (p<0.05, FC 1.5). (**C**) Gene ontology analysis of the TE downregulated and upregulated genes upon ADAT2 silencing (p < 0.04, gene count > 14). (**D**) Scatterplot of codon usage changes in the differentially translated genes (Down vs Up and Down-vs All other) in ADAT2 silenced cells. Dots in blue indicate ADAT-sensitive codons (codons read by ADAT-edited tRNAs). (**E**) Codon frequency of CGC, CCC, GCC, and ACC ADAT-sensitive codons comparing TE in upregulated, downregulated, and non-affected genes after ADAT2 silencing. * p < 0.05, ** p < 0.01, *** p < 0.001, ns = non-significant.

## Discussion

This study showed that the ADAT2/3 genes encoding the A-to-I tRNA editing complex are commonly amplified and/or overexpressed in cancer, and that ADAT2/3-depletion inhibits cancer cell growth hallmarks and tumorigenicity in human Liposarcoma cell lines. tRNA sequencing revealed decreased A-I editing of ANN tRNAs in ADAT2-deficient cells. It is worth mentioning that some tRNAs are more susceptible to ADAT knockdown than others likely reflecting the incomplete loss of ADAT2 editing activity in these LPS cells. Notably, Ribo-Seq revealed decreased translation of a set of mRNAs that are enriched in codons that depend on A-I editing for decoding. Thus, ADAT-mediated tRNA modification promotes oncogenesis by enhancing the translation of growth promoting genes that are enriched in NNC codons that lack cognate tRNAs and therefore require A-I tRNA editing for expression.

Although this study focused on human Liposarcoma, it is likely that the identified oncogenic function of ADAT2/3 is applicable to a broad range of different cancer types. Inspection of the DepMap dataset (https://depmap.org/portal) identified many different cancer cell lines that require ADAT2/3 for proliferation, in particular Rhabdomyosarcoma (ADAT2) and Diffuse Large B-cell Lymphoma (ADAT3)[24]. In further support of this widespread oncogenic role, a previous study found that ADAT2-depletion led to decreased proliferation of HeLa (cervical carcinoma) and HT-29 (colorectal carcinoma) cells[11]. Moreover, increased ADAT2 mRNA level is associated with poor prognosis in breast cancer patients and ADAT protein levels were found by immunohistochemistry staining to be higher in BRCA1 mutant compared with BRCA1 wild-type breast tumors[25], [26]. Taken together with our findings that ADAT2 and ADAT3 mRNA expression levels are elevated in most tumor types compared with the corresponding normal tissues we propose that ADAT2/3 is broadly relevant in cancer.

Our findings reported here add to the growing list of tRNA-modifying enzymes with important roles in cancer[27]. For example, other modifications in the anticodon[28], anticodon loop[29], as well as in the body of the tRNA can impact tRNA function and/or abundance[30], [31], [32], [33], [34], [35], [36]. Interestingly, the growth promoting and oncogenic influence of these different tRNA modifications converge mechanistically at the level of codon-biased mRNA translation whereby subsets of mRNAs linked to several different hallmarks of cancer including cell-cycle genes are enriched in certain codons and can therefore be coordinately controlled at the mRNA translational level through changes in the modification status of the corresponding tRNAs that are preferentially required for decoding these mRNA subsets[37], [38].

Enzymes involved in maintaining the epitranscriptome are emerging as promising new targets for cancer therapy[39], [40]. Pharmacological inhibition of the m^6^A methyltransferase (MTase) METTL3, represents a promising strategy for treating acute myeloid leukemia (AML) and likely other cancer types[41]. Our results uncover an oncogenic role of ADAT2/3-mediated tRNA editing and identify this adenosine deaminase as a possible new pharmacologically tractable target for cancer therapy. The available crystal structures of the ADAT2/3 complex[3], [4] as well as cryo-EM structures of the ADAT2/3 heterodimer bound to its substrate tRNA[5] should facilitate the development of small molecules inhibitors of this enzyme.

## Materials and methods

### Cell Lines

BJ human fibroblasts and HEK293T cells were purchased from ATCC. LPS6[42] was a gift from Eric Snyder, LPS141[42] and LPS853[43] were gifts from Jonathan Fletcher, and 93T449[44] was a gift from Florence Pedeutour. BJ and HEK293T were cultured in DMEM supplemented with 10% FBS and 1X penicillin/streptomycin. LPS141 and 93T449 cells were cultured in RPMI 1640 medium supplemented with 15% FBS and 1X penicillin/streptomycin. LPS853 cells were cultured in IMDM medium supplemented with 15% FBS and 1X penicillin/streptomycin. LP6 cells were cultured in DMEM/F12 medium supplemented with 10% FBS, 1% Glutamax and 1X penicillin/streptomycin. All cell lines were cultured in the presence of 5% CO2 at 37°C.

### Plasmid Construction and shRNAs

For overexpression experiments, the ADAT2 and ADAT3 full-length cDNAs were cloned in the pBabe-puro vector using BamHI and SalI restriction sites. A Flag-tag was added in the N-terminal region for both proteins. Mutant ADAT2 E73A was produced by directed mutagenesis using the Q5 Site-Directed Mutagenesis Kit (New England Biolabs). Mutagenesis primers are detailed in **Table S1**. For the ADAT2 rescue experiments, wild-type ADAT2, and the catalytic mutant (E73A) form were mutated to avoid shRNA silencing using the primers detailed in **Table S1** and using the same directed mutagenesis kit. For knockdown experiments, shADAT2 #1 (#653), shADAT2 #4 (#656), shADAT3 #4 (#576) and shADAT3 #5 (#577) were purchased from Sigma Aldrich.

### Copy Number Analysis

Genomic DNA was isolated using Quick-DNA microprep kit (Zymo) following the manufacturer’s instructions. ADAT2 and ADAT3 copy number alteration was evaluated using gene specific TaqMan Copy Number Assay (Thermo Fisher Scientific) according to the manufacturer’s instructions.

### Quantitative RT-PCR

For gene expression analysis, total RNA was isolated with Trizol Reagent (Invitrogen). Template cDNA synthesis was performed using the SuperScript III Reverse Transcriptase (Invitrogen). The levels of specific RNAs were measured by quantitative reverse transcription-PCR (qRT-PCR) using the StepOne real-time PCR machines and the Fast SybrGreen PCR mastermix (ThermoFisher, 4385612) according to the manufacturer’s instructions. All samples, including the template controls, were assayed in triplicate. The relative number of target transcripts was normalized to GAPDH or ACTIN. The relative quantification of target gene expression was performed with the standard curve or comparative cycle threshold (CT) method. All primers were purchased from Genewiz and are described in **Table S1**.

### Protein extraction and western blotting

Cells were lysed and proteins extracted with Passive Lysis Buffer (Promega) according to the manufacturer’s instructions. Protein concentration was measured using the Bradford method (Bio-Rad Laboratories). Samples were separated by sodium dodecyl sulfate– polyacrylamide gel electrophoresis (SDS-PAGE) Novex 4-20% Tris-Glycine gel (Thermo Fisher Scientific) and transferred to nitrocellulose membranes (Bio-Rad). Immunoreactive proteins were visualized by enhanced chemiluminescence (Thermo Fisher Scientific). The ADAT2 (ab135429), ADAT3 (ab247133), and Tubulin (ab6064) antibodies were obtained from Abcam. The Vinculin antibody (sc-73614) was purchased from Santa Cruz and β-Actin (8H10D10) (#3700) from Cell Signaling.

### Virus production and generation of stable knockdown and overexpression cells

Generation of stable knockdown and overexpression cells via virus transduction was performed described previously[45]. For knockdown experiments, the procedure involved co-transfecting shRNA-containing pLKO.1 vectors with pLP1, pLP2, and VSVG into HEK293T cells. For overexpression, Gag-Pol and VSVG plasmids were co-transfected with pBabe vectors containing either wild-type ADAT2, wild-type ADAT3, ADAT2 catalytic dead mutant (ADAT2 E73A), or an empty vector as a control. The viruses were collected at 48 h and 72 h after transfection and used to infect cells. Infected cells were selected using puromycin (2.5ug/mL) added to the culture medium 48 hr after infection. shRNA-expressing BJ fibroblast cells were maintained in medium supplemented with puromycin (2.5ug/mL).

### Proliferation and cell viability assays

Cell proliferation / cell viability was measured using the XTT metabolic assay (Cyman Chemical) and 2D colony formation using crystal violet staining. For XTT analysis, either 500 or 1,000 cells were seeded in 96-well plates and allowed to grow for 72 hours, after which dye reduction was recorded on a spectrophotometer at 450 nm. Crystal violet staining was performed by seeding 1,000 or 2,000 cells in each well of a 6-well plate. After 2-3 weeks, individual wells were fixed in 4% paraformaldehyde and stained with crystal violet. Pictures were obtained after extensive washing and drying, and the staining reagent was resolubilized in 1% acetic acid and quantified at 590 nm as an indirect measure of cell number.

### Invasion assay

Cell invasion was assayed in FluoroBlok cell culture inserts (Corning) coated on the upper side with Matrigel (Corning) mixed with DMEM at a final concentration of 300 μg/mL. 75,000 LPS853, cells or 50,000 LPS6 cells in DMEM containing 0.2% FBS were seeded in the upper chamber and 10% FBS was added to the bottom chamber as a chemoattractant. Cells were allowed to invade for 22 h at 37°C, 5% CO2 atmosphere. Cells were fixed in 100% methanol and stained with a 300 nM DAPI solution. Images were obtained using a florescence microscope and cells counted using Image J.

### Soft agar colony formation assays

5,000 LPS853 or LPS6 cells were mixed with 0.35% top-agar (SeaPlaque, Lonza) and were plated onto 0.7% base-agar (SeaPlaque, Lonza) in six-well plates. Three weeks (LPS853) or five weeks (LPS6) after plating the cells into soft agar, the plates were stained with violet crystal and imaged using a EVOS FL auto plate imager (Thermo Fisher Scientific) at 2x (LPS853) or 4x (LPS6). Colony numbers were counted using image J.

### In vivo studies

Female NU/J (Nude) immunodeficient mice (Jackson Laboratory) aged 4-6 weeks were used for subcutaneous injections to investigate the role of ADAT2 in tumor formation. For this purpose, 5×10^5^ LPS853 cells with stable knockdown of ADAT2 (shADAT2 #1 and #4) or a negative control (shGFP) were used. The indicated number of cells were mixed with serum-free medium and growth factor reduced Matrigel (Corning) (1:1) and injected into the right flank of 6-8 mice for each condition. Tumor growth was monitored twice a week using calipers. The tumor volume was calculated using the formula 1/2(length × width^2^). End-point tumors were collected and photographed.

### Small RNA sequencing

To analyze the tRNA levels and editing, small RNA sequencing was performed. Total RNA was isolated with Trizol Reagent (Invitrogen) and treated with recombinant AlkB and AlkB D135S for demethylation for 2 h as detailed in[46]. Demethylated small RNA was purified using the RNA Clean and Concentrator –5 kit (Zymo Research) and the library prepared using the NEBnext Multiplex small RNA library prep for Illumina (New England Biolabs) following the manufacturer’s instructions. Amplified libraries were selected and purified by gel-extraction on 6% native TBE gels (Thermo Fisher Scientific). Libraries were sequenced with Illumina NextSeq 500.

### Ribosome footprinting (Ribo-seq)

LPS853 cells stably silenced for ADAT2 (shADAT2), or control shRNA (shGFP) were grown to 80-90% confluence to obtain at least 50 million cells per condition and replicate(n=2). Ribosome footprinting was performed according to TruSeq® Ribo Profile system (Illumina) with some modifications described in[30]. For library preparation the NEBNext Multiplex Small RNA Library Prep set for Illumina (New England Biolabs) was used. In parallel, total RNA input samples were isolated, and fragmented and the library obtained using the TruSeq stranded total RNA Kit (Illumina). Amplified libraries were selected and purified by gel-extraction on 6% native TBE gels (Thermo Fisher Scientific). Libraries were sequenced with Illumina NovaSeq SE50. To perform the analysis, the sequences of both input and Ribo-seq samples were processed to obtain clean reads by trimming the adapters and filtering out low-quality sequences. For the Ribo-seq input data, the clean reads were aligned to reference genome sequences using STAR[47], resulting in BAM mapping files. HTSeq[48] was then used to calculate the read counts for each gene from GENCODE gene mode. For the cleaned Ribo-seq data, ribosome-protected fragments (RPFs) were collapsed into FASTA format using fq2collapedFa. Codon occupancy analysis and translation efficiency analysis were performed using RiboToolkit (https://bioinformatics.caf.ac.cn/RiboToolkit_demo)[49]. Translation efficiency was calculated by dividing RPF abundance on CDS by its mRNA abundance of input sample. Differential translation genes were defined by a p-value <0.05 and a 1.5-fold change. **Supplemental Dataset 1**.

### Functional annotation of candidate genes

The genes obtained after the Ribo-seq (genes with up-or down-translation efficiency) were processed by The Database for Annotation, Visualization and Integrated Discovery (DAVID) (https://david.ncifcrf.gov) for functional annotation.

### TCGA data analysis

To study the frequency of genomic alterations in samples from the TGCA the cBioportal (www.cbioportal.org) was used. The mRNA expression levels were obtained from Firebrowse (www.firebrowse.org).

### Statistical analysis

Quantification and statistical analysis methods were described in individual method sections and Figure legends. Results are expressed as the mean ± standard deviation of three different experiments (n=3) unless specified. Statistical significance was determined by Student’s t-test analysis (two-tailed) and differences were considered significant at a p-value < 0.05.

### Data availability

High-throughput sequencing data have been deposited in the Gene Expression Omnibus (GEO) under the accession numbers GSE234132 (tRNA-seq) and GEO Pending (Ribo-seq).

### Code availability

The software and algorithms for data analyses used in this study are all well-established from previous work and are referenced throughout the manuscript.

### Statistics and reproducibility

Detailed statistical analysis methods and sample numbers (n) were described in individual figure legends. GraphPad Prism and Microsoft Excel were used for data presentation. Paired or unpaired two-tailed Student t-tests, Fisher’s exact tests, or Mann-Whitney two-sided U tests were used for two comparisons. Statistical significance is considered for all analysis where *p<0.05, **p<0.01, ***p<0.001, ****p<0.0001.

## Acknowledgments

R.I.G. was supported by an Outstanding Investigator Award (R35CA232115) from the National Cancer Institute (NCI) of the NIH.

## Author contributions

J.R-M and R.I.G. designed the research. J.R-M. performed most of the experiments. Q.L. and X.X performed all the bioinformatics analysis. R.A. performed the mouse xenograft experiments under A.G. supervision. J.R-M, T.R.A., and R.I.G. analyzed the data and wrote the paper with input from other authors.

