## Supplementary Information for "ADAT2/3-mediated tRNA editing promotes cancer cell growth and tumorigenicity"

**Contents**

Supplementary Figure 1
Supplementary Figure 2
Supplementary Figure 3
Supplementary Figure 4
Supplementary Figure 5
Supplementary Table 1

**A**

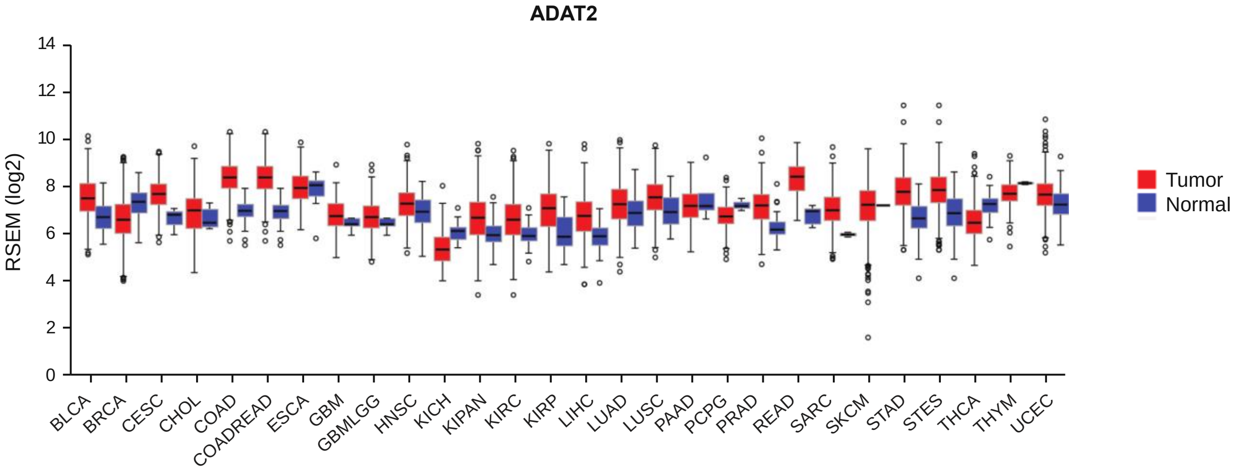


**B**

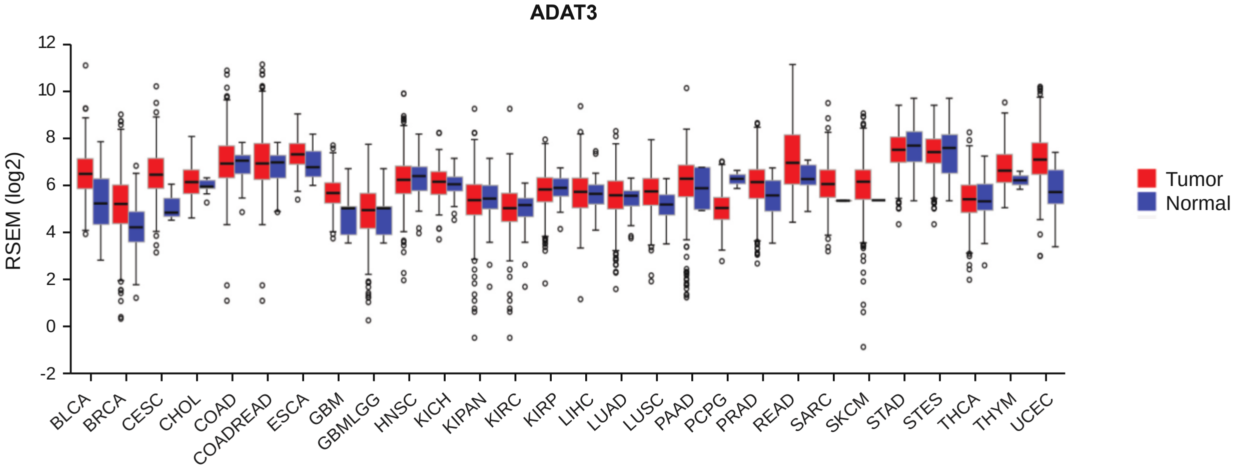


**Figure S1. ADAT2 and ADAT3 are overexpressed in most tumor types** (**A**) ADAT2 mRNA levels in tumors and normal samples represented in the TCGA. (**B**) ADAT3 mRNA levels in tumors and normal tissues. BLCA: Bladder Urothelial Carcinoma, BRCA: Breast invasive carcinoma, CESC: Cervical squamous cell carcinoma and endocervical adenocarcinoma, CHOL: Cholangiocarcinoma, COAD: Colon adenocarcinoma, COADREAD: Colorectal adenocarcinoma, ESCA: Esophageal carcinoma, GBM: Glioblastoma multiforme, GBMLGG: Glioma, HNSC: Head and Neck squamous cell carcinoma, KICH: Kidney Chromophobe, KIPAN: Pan-kidney cohort (KICH+KIRC+KIRP), KIRC: Kidney renal clear cell carcinoma, KIRP: Kidney renal papillary cell carcinoma, LIHC: Liver hepatocellular carcinoma, LUAD: Lung adenocarcinoma, LUSC: Lung squamous cell carcinoma, PAAD: Pancreatic adenocarcinoma, PCPG: Pheochromocytoma and Paraganglioma, PRAD: Prostate adenocarcinoma, READ: Rectum adenocarcinoma, SARC: Sarcoma, SKCM: Skin Cutaneous Melanoma, STAD: Stomach adenocarcinoma, STES: Stomach and Esophageal carcinoma, THCA: Thyroid carcinoma, THYM: Thymoma, UCEC: Uterine Corpus Endometrial Carcinoma.

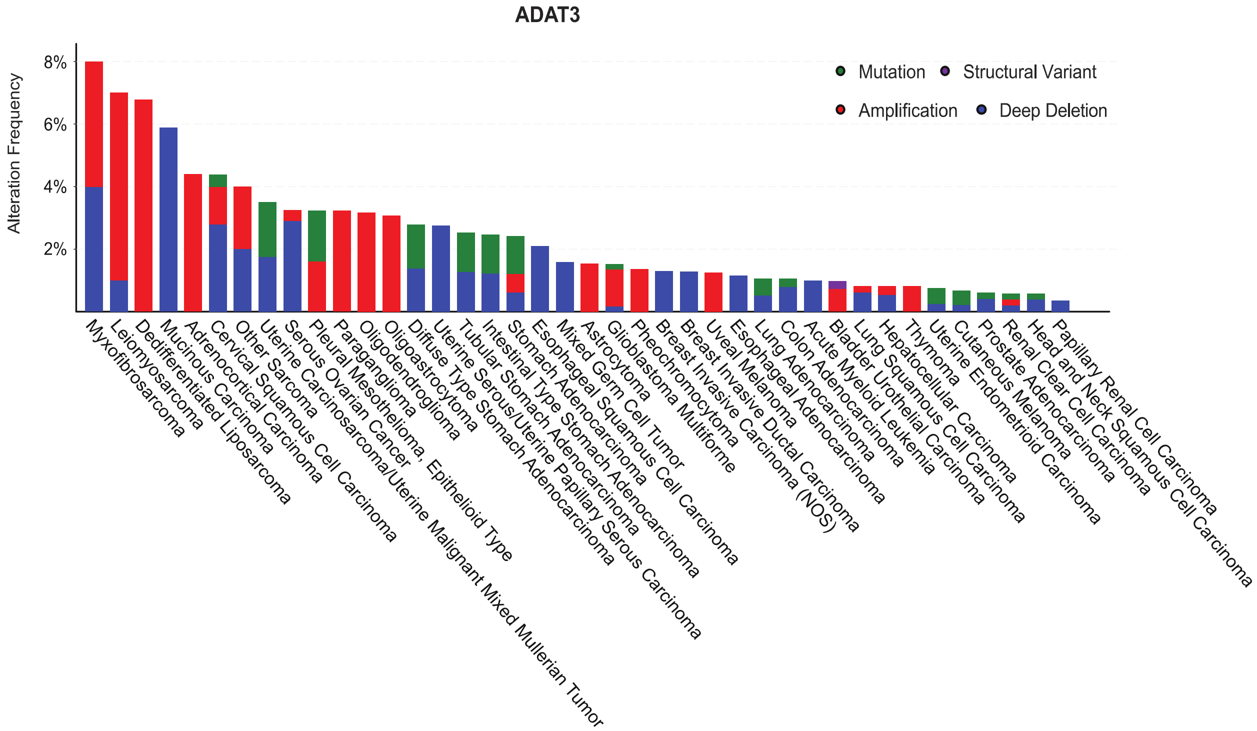


**Figure S2. *ADAT3* is amplified in liposarcoma**. (**A**) *ADAT3* genetic alterations (mutations, amplifications, deletions, and structural variants) in TCGA tumors. “Other sarcomas” refers to Undifferentiated Pleomorphic Sarcoma/Malignant Fibrous Histiocytoma/High-Grade Spindle Cell Sarcoma.

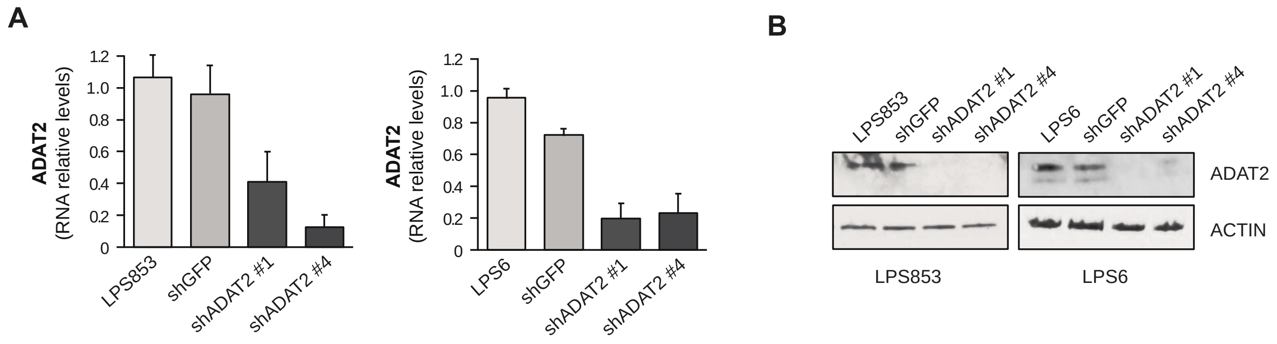


**Figure S3. ADAT2 knockdown in liposarcoma cells**. ADAT2 was silenced in LPS853 and LPS6 cell lines using two shRNAs (shADAT2 #1 and shADAT2 #4). shGFP and wild-type cells were used as negative controls. (**A**) ADAT2 mRNA relative levels assayed by qPCR. Error bars indicate standard deviations (n=3). (**B**) Representative Western Blotting for ADAT2. Actin was used as loading control.

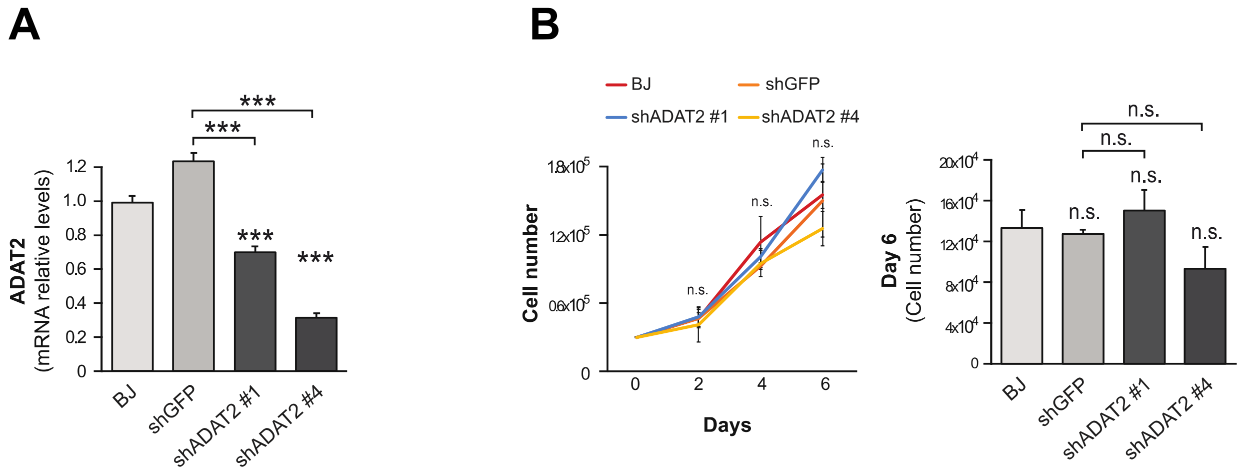


**Figure S4. ADAT2 depletion in non-transformed fibroblasts.** ADAT2 knockdown in BJ human fibroblasts using two shRNAs (shADAT2 #1 and shADAT2 #4). shGFP and wild-type cells were used as a negative control. (**A**) ADAT2 relative mRNA levels assayed by qPCR in BJ cells (n=3). (**B**) Proliferation assay in BJ cells (n=3). Error bars indicate standard deviations. Asterisks denote statistical significance assessed with Student's t-test (two-tailed). *** $p < 0.001$, ns = non-significant.

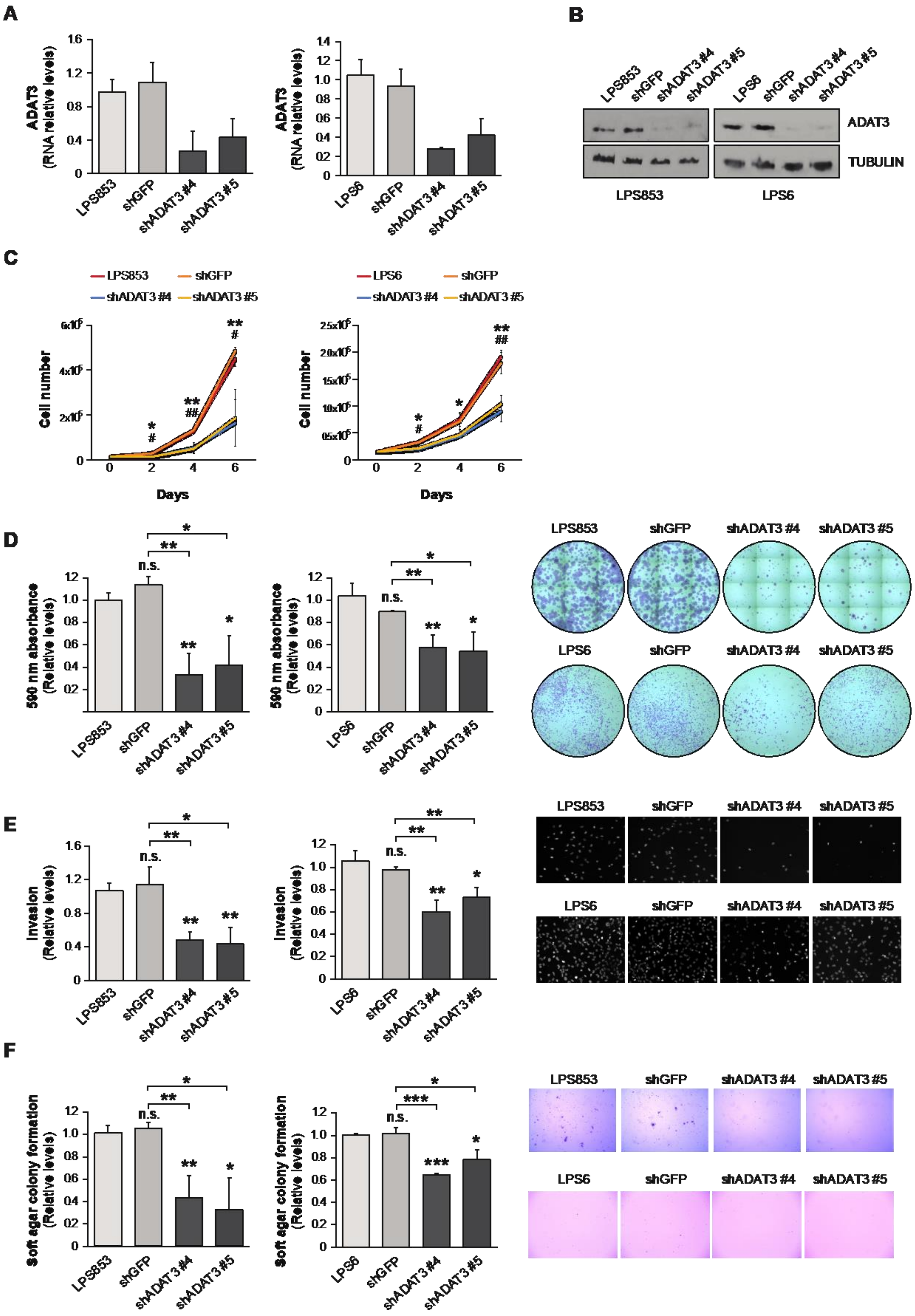
A
ADAT3 (RNA relative levels)
LPS853
shGFP
shADAT3 #4
shADAT3 #5
LPS6
B
ADAT3
TUBULIN
LPS853
LPS6
C
Cell number
Days
D
590 nm absorbance (Relative levels)
n.s.
E
Invasion (Relative levels)
F
Soft agar colony formation (Relative levels)

**Figure S5. ADAT3 knockdown decreases oncogenic hallmarks in liposarcoma cells**. ADAT3 knockdown using two shRNAs (shADAT3 #4 and shADAT3 #5) in LPS853 and LPS6 cells. shGFP and wild-type cells were used as a control. (**A**) ADAT3 mRNA relative levels assayed by qPCR (n=3). (**B**) Representative Western Blotting for ADAT3 protein. Tubulin was used as loading control. (**C**) Proliferation assay (n=3). (**D**) 2D colony formation assay. Quantification (left) and representative images (right) (n=3). (**E**) Invasion assay. Quantification (left) and representative images (right) (n=3). (**F**) 3D soft agar colony formation assay. Quantification (left) and representative images (right) (n=3). Error bars indicate standard deviations. Asterisks denote statistical significance assessed with Student's t-test (two-tailed). * $p < 0.05$, ** $p < 0.01$, *** $p < 0.001$, ns = non-significant.

### Supplementary Table 1

qPCR primers:

| **Primer** | **Sequence** |
|---|---|
| ADAT2 _F | ACCAAAAATGCTACTCGACATGC |
| ADAT2 _R | ACCAGCGGGATTTTCATCAGG |
| ADAT3 _F | CCGGATCCCTCAGGGGTTA |
| ADAT3_R | GAGACAGAGACGGGAGCAGA |
| GAPDH_F | TGCACCACCAACTGCTTAGC |
| GAPDH_R | GGCATGGACTGTGGTCATGAG |
| ACTIN_F | ACAGAGCCTCGCCTTTG |
| ACTIN_R | CCTTGCACATGCCGGAG |

Mutagenesis primers:

| **Primer** | **Sequence** |
|---|---|
| ADAT2_E73A_F | CGACATGCAGCAATGGTGGCC |
| ADAT2_E73A_R | AGTAGCATTTTTGGTTTGGTTAAC |
| ADAT2_NoSh_F | ACGGATGCCAGAATGAACGATTTGGTGGTTG |
| ADAT2_NoSh_R | ACACGACTAATGGGATTTTCATCAGGCGG |
| ADAT3_NoSh_F | TATTGTGTTTGGCTGGGCCGGCCTCGGG |
| ADAT3_NoSh_R | ACATTTCTAGTGCGTGGGGGCTGCCGGCATC |
